# Keys to the cabinet: unlocking biodiversity data in public entomology collections

**DOI:** 10.1101/2024.05.10.593424

**Authors:** Joel F. Gibson, Mackenzie Howse, Claire Paillard, Cassandra Penfold, Alannah Penno, Genevieve E. van der Voort, Dezene P.W. Huber

## Abstract

Canadian entomology collections contain valuable biodiversity and ecological data, but they must be digitized in order to be usable by those working outside of the collections. Multiple analyses of the digital database of the Odonata collection at the Royal British Columbia Museum were conducted. These analyses reveal that complete digital datasets can be used to explore questions of historical and current geographical distribution and species composition differences based on ecoprovince and elevation. The results of these analyses can be used directly in conservation and climate change impact mitigation decisions. These analyses are only possible because the Odonata collection has received concerted effort to digitize all specimen records. The full value of long-term historical insect biodiversity data can only be accessed once collections are digitized. Additional training and employment of collection management and curatorial staff is essential to optimize the use of abundant, but underutilized, Canadian biodiversity data.

## Introduction

Entomology collections across Canada are a valuable source of ecological and biodiversity data. Each collection houses, maintains, and makes available preserved and labeled terrestrial arthropod specimens. While physical specimens - on pins, in ethanol, in envelopes, and on slides - are the core material of collections, it is the data represented by those specimens that are of immeasurable value to those working within and outside of the collections themselves. The digital age has allowed accelerated capture of the physical data of collections in electronic formats that can be shared around the globe.

There is general acknowledgement that digitized, publicly available biodiversity records are of immense value for researchers working outside of natural history collections (*e.g*., Cardoso *et al*. 2011; Nelson and Ellis 2018; Miller *et al*. 2020). Large datasets from entomology collections are regularly used for meta-analyses. For instance, species distributions and historical changes in those geographical ranges can be observed using regionally focused entomological collection datasets (Favret and DeWalt 2002). Consulting entomology collection records usually adds records of species not previously known to be present within a geographical region. For example, by digitizing the Illinois Natural History Survey (INHS) collection, Favret and DeWalt (2002) added four new species records to the relatively well-known species list of Ephemeroptera of Illinois. Conventional entomology collection label data (species identification, locality, and date) can be easily combined with other datasets including physiological data, historical climactic data, land use data, genomic data, and population genetics studies. The long timescale of entomology collection data can be a useful historical complement to new studies that focus on intense monitoring of a target species in a limited geographical area (Kharouba *et al*. 2018). In some cases, original metaanalyses of historical entomological data take on a focus that was likely inconceivable to the collectors of the original material. For example, Kharouba *et al*. (2018) completed a comprehensive review of studies that employed entomology collection data to investigate historical patterns of climate change impacts.

The unfortunate reality is that most entomology electronic databases lack substantial amounts of the collection’s specimen data. In a recent estimate, only 7% of specimen data in Canadian entomology collections has been digitally captured (Cobb *et al*. 2019). Even the data that have been digitized may not be usable without improvement or completion. Digital specimen records that do not contain a full complement of metadata (locality, georeferenced coordinates, collection date, collector, species identification) are considered ‘skeletal’ or incomplete, although in some instances, records containing lower data resolution (*e.g.*, family level identifications, localities without coordinates) can still be used for limited analyses.

Taxonomic inconsistencies and geospatial data anomalies are the two main sources of error in digitized biodiversity datasets (Nelson and Ellis 2018). Updating out-of-date taxonomic terms and standardizing locality coordinates is a time-consuming process that is often necessary with older specimen records (Giberson and Burian 2017). Even when data anomalies have been addressed, entomological collection data are often unevenly distributed temporally and spatially (Kharouba *et al*. 2018). Entomological specimens collected over decades or even centuries by sometimes hundreds of collectors were not collected according to a predetermined pattern or design. One challenge of analysing digital entomology datasets is knowing the degree of geographical and temporal specialization or biases in a dataset.

In this study we interrogated a single, publicly available entomological collection of dragonflies and damselflies (Odonata) collected between 1913 and 2021, in British Columbia. We performed ecological, spatial, and taxonomic analyses to examine biological, temporal, spatial, and methodological patterns. While some completion was performed and inconsistencies were corrected, no additional collecting or curatorial efforts were made in this study. Our study provides new insights into the diversity of British Columbia’s Odonata and demonstrates the growing potential for entomologists and others to use and improve existing digitized records for biodiversity and conservation research.

## Material and Methods

The core dataset for this study was the Odonata collection at the Royal British Columbia Museum (RBCM). The RBCM Entomology Collection is considered a small, tier-three collection (100,000 to 1,000,000 specimens) according to the categorization of Cobb *et al*. (2019). The biodiversity of British Columbia’s Odonata was a targeted research and collections goal of the museum for many years, led by the curator, Dr. Rob Cannings (Cannings 2023). Previous studies (Cannings 2019; Cerini *et al*. 2021; Cannings 2023) have made use of portions of the dataset.

All Odonata database records were downloaded from the RBCM database on February 7, 2023. The dataset was trimmed to include only records that were confirmed to have been collected within British Columbia.

Database completeness was assessed using the degree of detail included for each record for collection location, collection date, and taxonomic identification. Overall data completeness was visualized using a Sankey diagram using the R package *ggsankey* (Sjoberg 2021). The list of species names in the database was compared to the World Odonata List in the Catalogue of Life (Paulson et al. 2023). Current conservation status for each species was derived from the October 2022 version of the Committee on the Status of Endangered Wildlife in Canada (COSEWIC) list (BC C.D.C. 2023). A heat map of Odonata specimen occurrences was produced using ArcGIS Pro 3.2.1. Coordinates were geographically limited to British Columbia with a screen unit point radius of 10, locked to a scale of 1:5,000,000. To provide a point of reference for where specimens were collected, an overlay of major highways was included. Habitat descriptions were extracted from all records and then manually assigned to categories. These data were then incorporated into a Venn diagram using the R packages *ggVenn* (Yan 2023) and *ggVenndiagram* (Gao et al. 2021). Locations of specimen entries containing valid latitude and longitude coordinates, but not elevation records, were entered into Google Earth Pro (7.3.6.9345, 2022) and elevation values were retrieved. Odonata species similarity between elevation ranges was calculated using the faunal similarity index (FS=NC/(N1+N2-NC), NC=number of species in common, N1 and N2 = number of species collected in each elevation range) (Kearns 1992).

## Results

We recovered a total of 34,687 specimen records of Odonata collected in British Columbia. Collection dates ranged from 1913 to 2021. A complete list of specimens with data are provided in Supplementary Material. Completeness of the dataset was evaluated in various ways (Figure 1).

**Figure 1.**
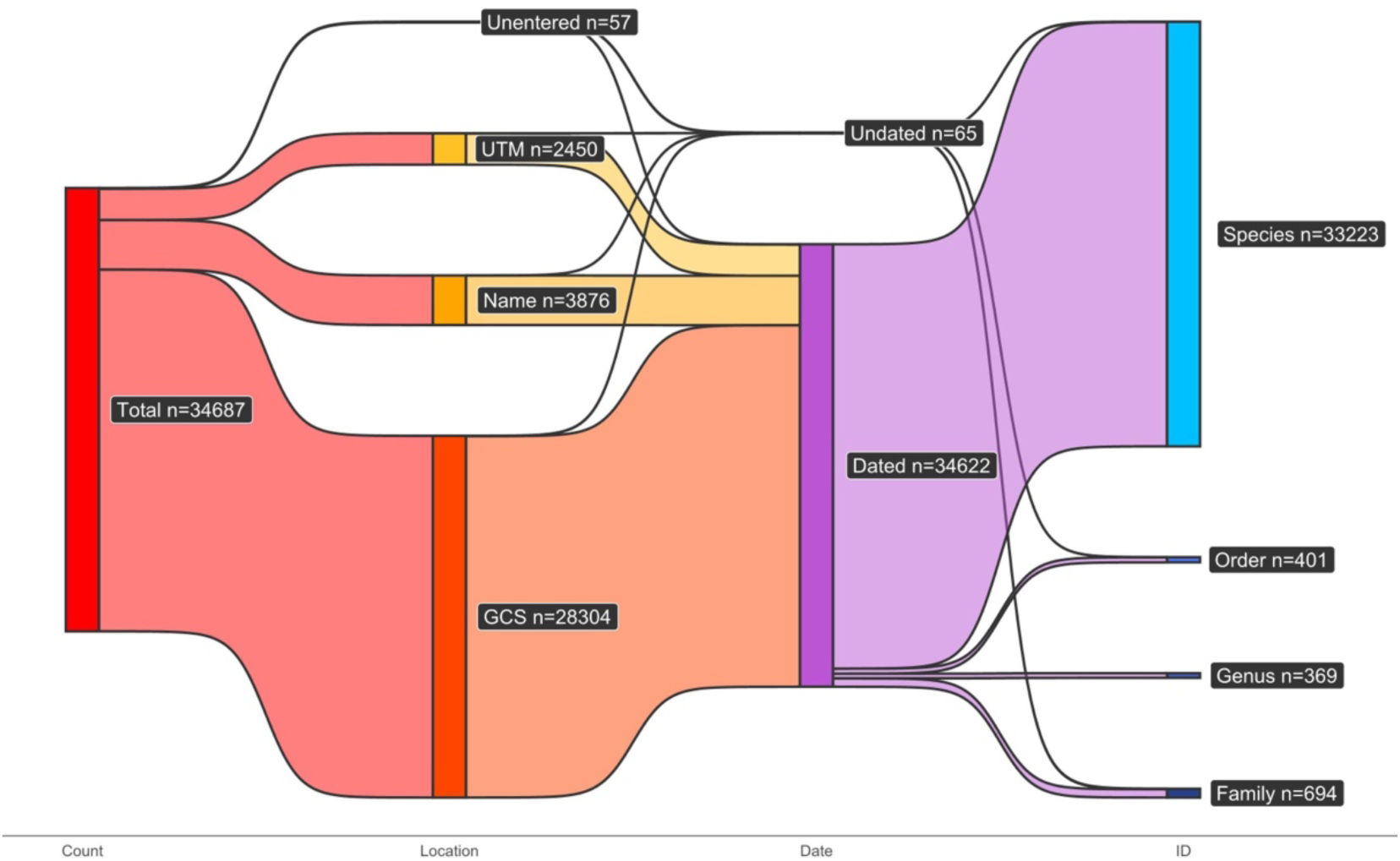
Sankey diagram of RBCM British Columbian Odonata specimen database entry completeness. Figure created in R version 4.2.3 with the package *ggsankey* (Sjoberg 2023).

For location data completeness, the degree of geographic accuracy was ranked. Full latitude and longitude coordinates were considered greater than UTM coordinates, which was considered greater than a locality name alone, which was considered greater than a blank location entry. Of the 34,687 records, 28,304 (81.6%) had full latitude and longitude data, 2,450 (7.1%) had UTM data, 3,876 (11.2%) had a locality name, and 57 (0.2%) had no location data beyond “British Columbia.”

For taxonomic completeness, entries were ranked by phylogenetic level. Of the 34,687 records, 33,223 (95.8%) had species-level identifications, 369 (1.1%) had genus identifications, 694 (2.0%) had family identifications, and 401 (1.2%) were identified only to order as Odonata. Of the 34,687 records, 34,622 (99.8%) had a collection date recorded.

On a per family basis (Table 1), species-level identification ranged from 95.4% of records (Gomphidae) up to 100% of records (Cordulegastridae, Macromiidae, Calopterygidae, Petaluridae). Collection date was recorded for more than 99% of records for all families. Complete latitude and longitude data were recorded for as little as 52.5% of records (Cordulegastridae) to as much as 100% of records (Petaluridae).

**Table 1.** Summary table of data completeness for RBCM British Columbian Odonata specimens. Values represent percentage of specimens having data present in the database for each Odonata family.

| Family | Total | Genus<br>ID | Species<br>ID | Collection<br>Date | Latitude<br>Longitude |
| --- | --- | --- | --- | --- | --- |
| Coenagrionidae | 9217 | 96.3 | 96.1 | 99.6 | 82.8 |
| Libellulidae | 8829 | 99.6 | 99.2 | 100.0 | 79.6 |
| Aeshnidae | 8397 | 96.9 | 93.6 | 99.8 | 83.1 |
| Lestidae | 3678 | 99.6 | 99.3 | 99.9 | 81.0 |
| Corduliidae | 3642 | 99.0 | 98.7 | 99.9 | 81.7 |
| Gomphidae | 372 | 98.9 | 95.4 | 100.0 | 75.5 |
| Cordulegastridae | 99 | 100.0 | 100.0 | 100.0 | 52.5 |
| Macromiidae | 35 | 100.0 | 100.0 | 100.0 | 77.1 |
| Calopterygidae | 13 | 100.0 | 100.0 | 100.0 | 92.3 |
| Petaluridae | 4 | 100.0 | 100.0 | 100.0 | 100.0 |

One historic taxonomic name in the dataset, *Sympetrum occidentale* Bartenev 1915, needs to be updated to the current accepted name *Sympetrum semicinctum* (Say 1839) (Pilgrim and Von Dohlen 2007). One species, *Enallagma exsulans* (Hagen 1861), was recorded in the dataset, but is believed to only range as far west as Manitoba and Texas. The 27 records of this species are likely to be misidentifications or mislabelled locations. Up until 1934, there were fewer than 50 species names recorded for British Columbia in the database. By 2001, the number of species in the database plateaued at 85 species. Three species on Cannings’ (2023b) Odonata of BC list [*Archilestes californicus* McLachlan 1895, *Enallagma civile* (Hagen 1861), and *Pantala hymenaea* (Say 1839)] are not present in the collection, but were reported as present in British Columbia in other public sources (Cannings 1988; Cannings and Pym 2017; Lee and Cannings 2024). A total of 16 species had no new records in the database over the past 20 years. For one species, *Enallagma clausum* Morse 1895, the most recent record was July, 1997. Of the nine British Columbia Odonata species currently listed as Threatened or Endangered on the COSEWIC list (BC C.D.C 2023), six have not had any new records added since 2008 (Table 2).

**Table 2.** Odonata species of British Columbia currently listed by the Committee on the Status of Endangered Wildlife in Canada (COSEWIC) and the number of specimens held by RBCM, and date range of held specimens, and date of COSEWIC status assessments.

| Scientific Name | Common Name | # specimens in RBCM | Oldest RBCM record | Most recent RBCM record | Date of recent COSEWIC assessment |
| --- | --- | --- | --- | --- | --- |
| *Argia vivida* | Vivid dancer | 127 | 1980-06-30 | 2015-07-24 | May 2015 |
| *Calopteryx aequabilis* | River jewelwing | 11 | 1999-07-19 | 2014-06-24 | - |
| *Enallagma civile* | Familiar bluet | 0 | - | - | - |
| *Enallagma clausum* | Alkali bluet | 62 | 1938-07-15 | 1997-07-25 | - |
| *Ischnura damula* | Plains forktail | 40 | 1980-05-24 | 2003-07-28 | - |
| *Octogomphus specularis* | Grappletail | 14 | 1936-08-12 | 2000-07-31 | May 2021 |
| *Somatochlora brevicincta* | Quebec emerald | 24 | 2000-08-12 | 2001-08-10 | - |
| *Somatochlora forcipata* | Forcipate emerald | 20 | 1998-08-03 | 2002-07-24 | - |
| *Somatochlora kennedyi* | Kennedy's emerald | 22 | 2001-07-08 | 2004-07-22 | - |
| *Stylurus olivaceus* | Olive clubtail | 19 | 1913-06-11 | 2008-08-25 | May 2011 |

The number of records added to the database per decade greatly increased beginning in 1973 and continued to increase until a peak in 1993-2002 (Figure 2). This period coincided with the concerted effort led by Dr. Rob Cannings with substantial participation of others to document Odonata biodiversity in the province. While 177 different collectors contributed specimens to the database, all of the species level identification was completed by only 41 individuals. In fact, nearly 90% of the species identifications were completed by just five individuals - Rob Cannings, Gord Hutchings, Richard Cannings, Syd Cannings, and Leah Ramsey (Figure 3).

**Figure 2.**
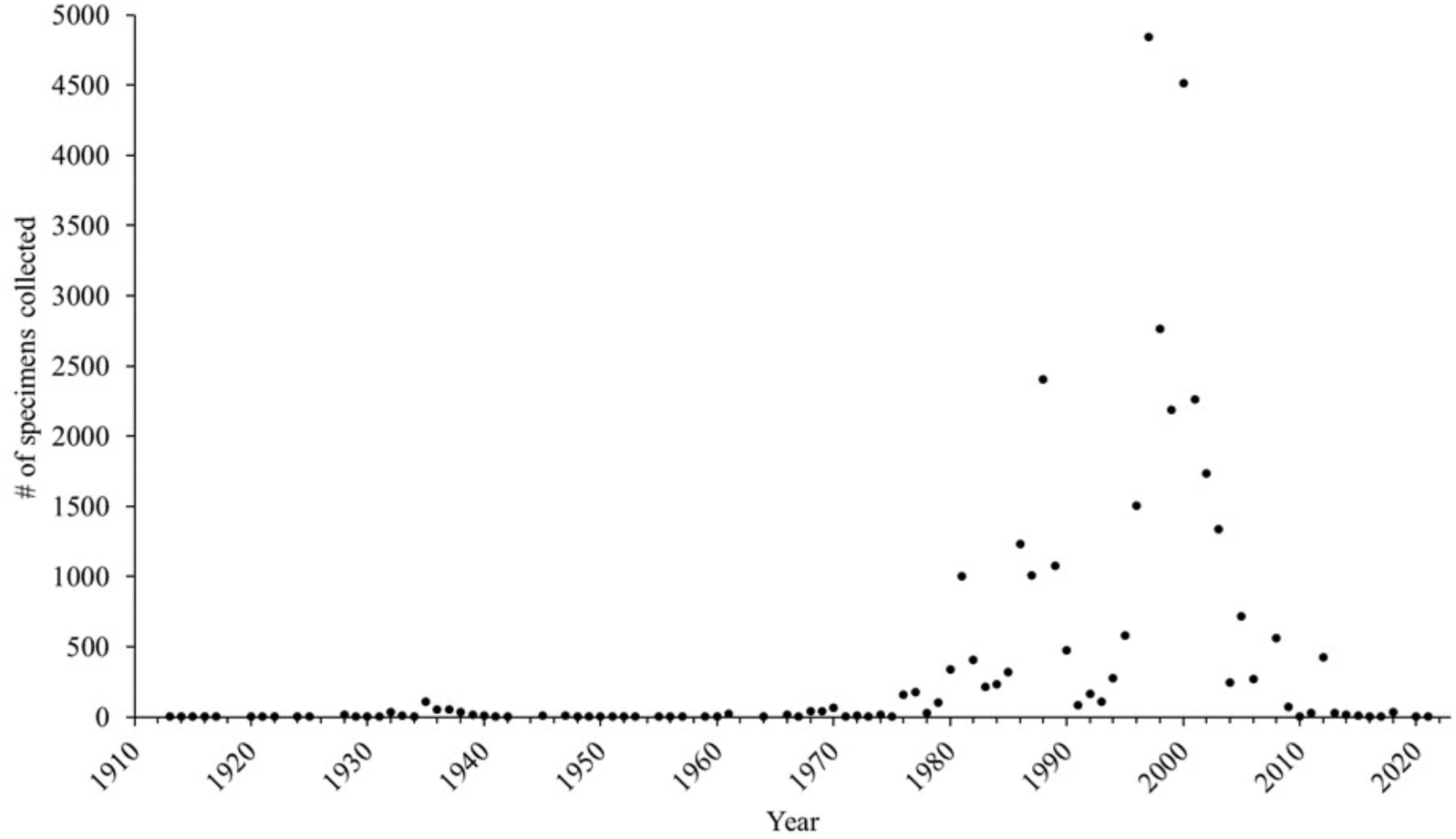
Number of British Columbia Odonata specimens collected by all collectors per year in the RBCM collection. 65 records without dates were excluded, total number of records = 34,622.

**Figure 3.**
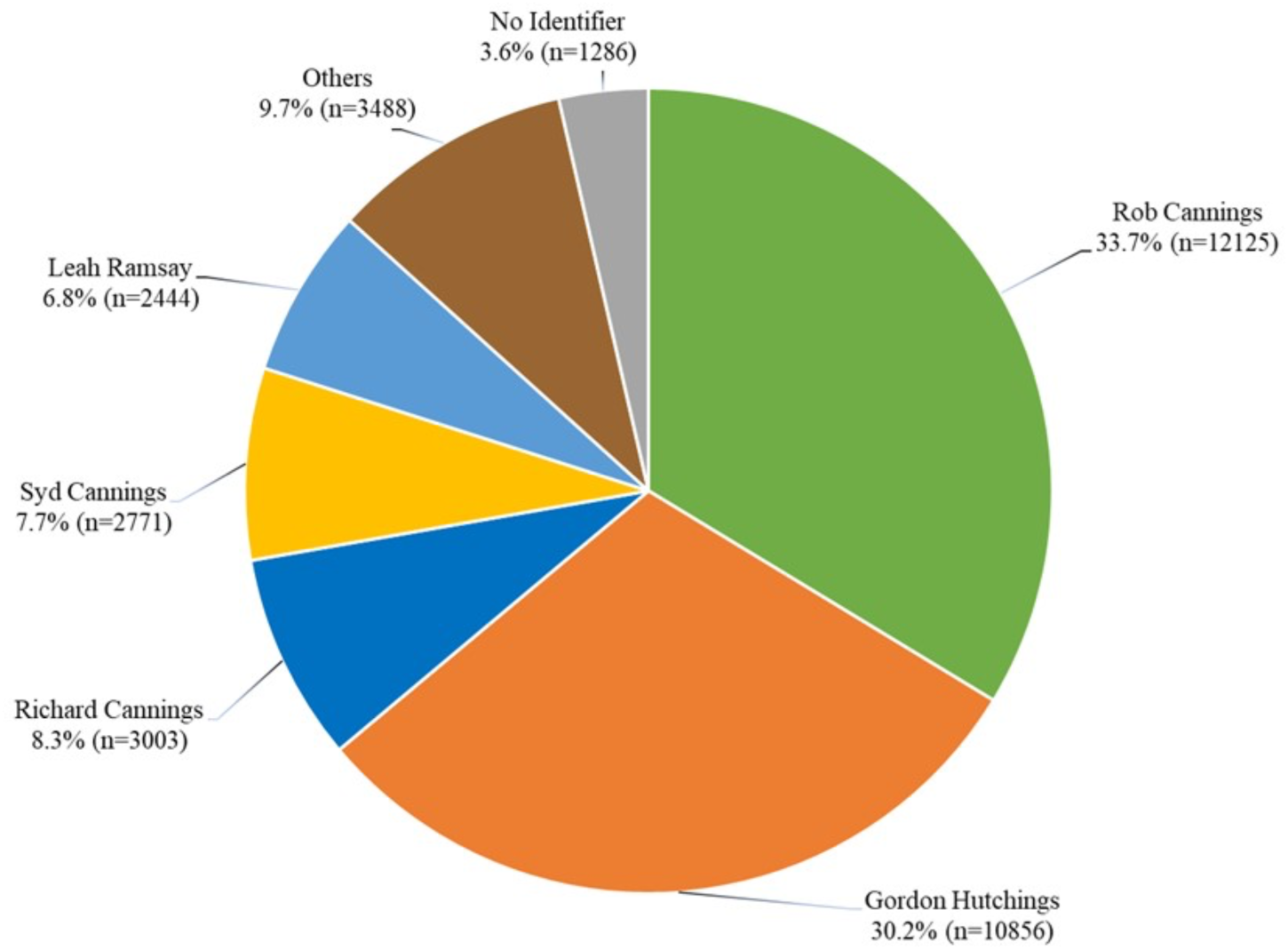
Identifiers of British Columbia Odonata specimens in the RBCM collection (n=34,687).

When all database records with verified latitude and longitude data (28,239 records) were placed on a single map (Figure 4), several patterns were observed. Major collecting hotspots were southern Vancouver Island (including Victoria), the southwest mainland (including Vancouver), and the southern Okanagan valley (including Penticton and Osoyoos). Outside of these areas, almost all records were along or near either Highway 97 or Highway 16. There were very few records from north of 56° N latitude. There were also very few records between the central coastal region east to Highway 97. Some targeted collection events were evident in the data, including a major collecting effort on Brooks Peninsula in 1981 (Cannings and Cannings 1983).

**Figure 4.**
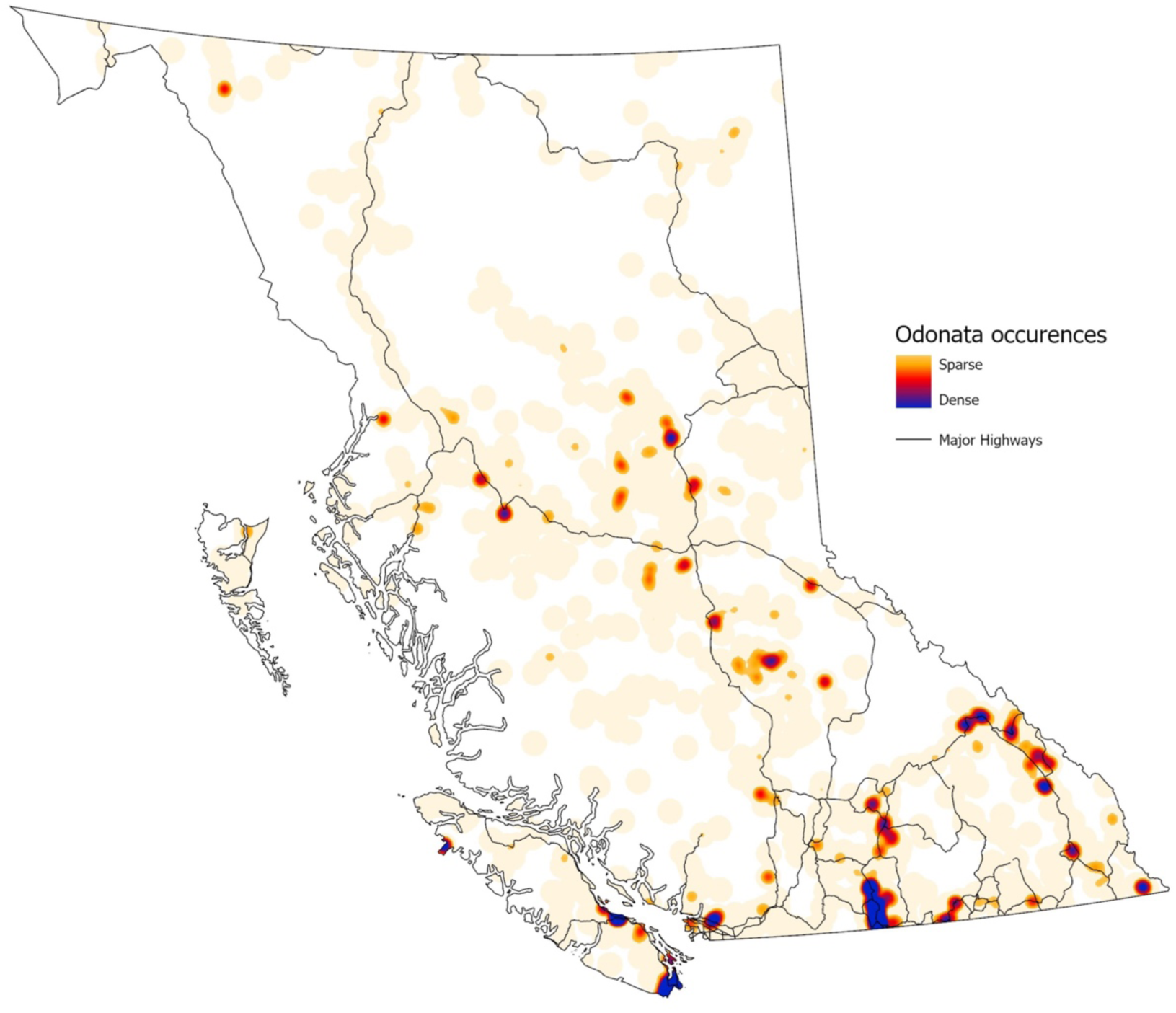
Heat map of RBCM British Columbia Odonata collection records in relation to major highways throughout British Columbia. Map generated with ArcGIS Pro 3.2.1.

Grouping collecting localities according to ecoprovinces (Demarchi 2011) allowed further geographic analysis (Figure 5). The number of records per ecoprovince varied, with Southern Interior Mountains having the most records (6,110) and, not including the small Southern Alaska Region, the Taiga Plains had the fewest (510). Species richness per ecoprovince also varied, with Southern Interior having the most recorded species (69) and Northern Boreal Mountains and Taiga Plains the fewest (33 each). An analysis based on similarity of species composition using only presence and absence data of each species produced a dendrogram showing overall similarities and differences between nine of eleven of British Columbia’s ecoprovinces (Figure 6). The Southern Interior Mountains, Southern Interior, Coast and Mountains, and Georgia Depression together formed one clade and the other northern and interior ecoprovinces form a second clade. This division of the province into two large metaregions aligns with the conclusions of Cerini *et al*. (2021) who used similar data but a different analytical approach.

**Figure 5.**
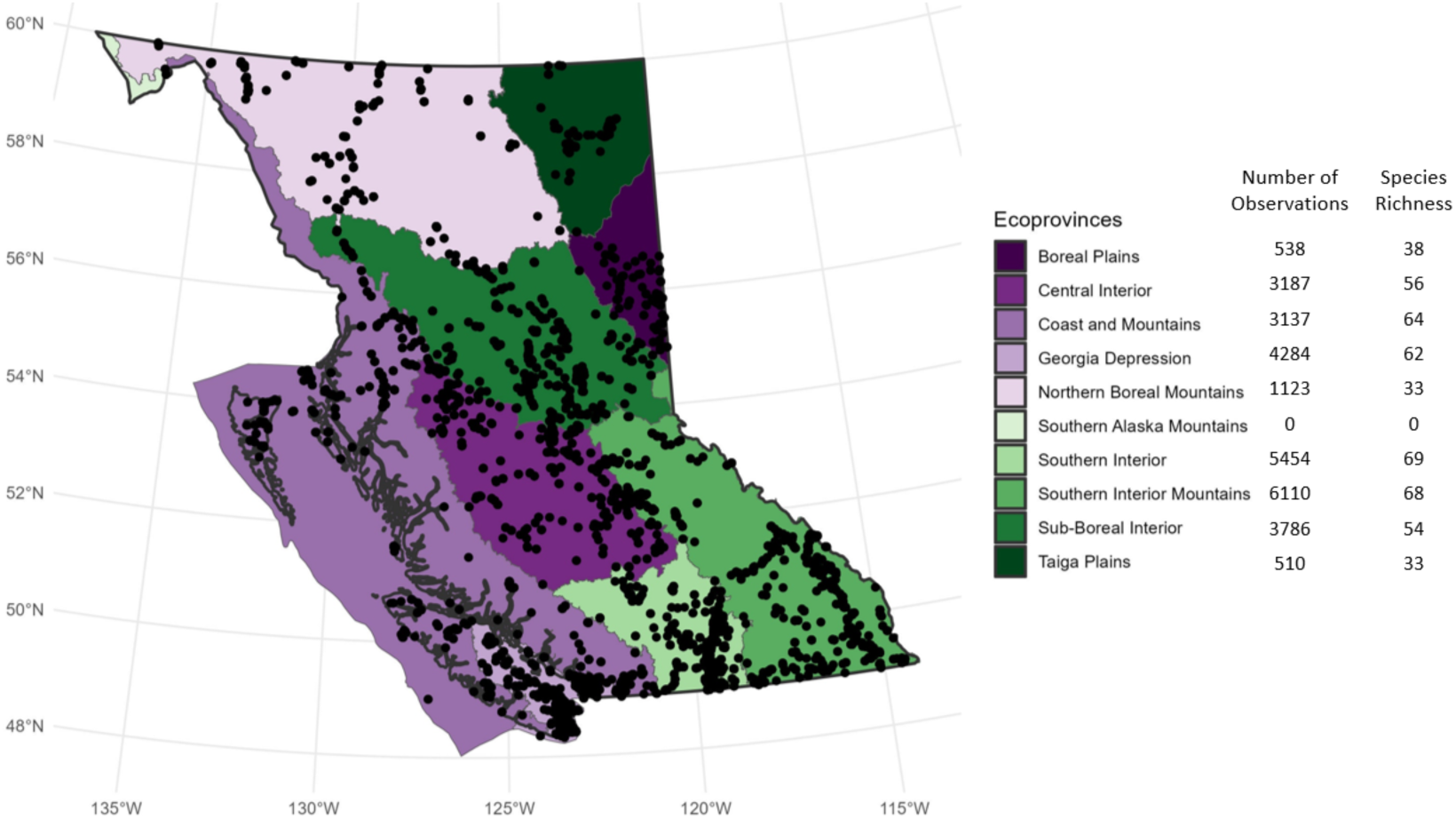
28,129 records of Odonata from the RBCM Entomology collection placed onto a map of British Columbia including ecoprovince designations. Map generated using R version 4.2.2 and the *bcmaps*, *sf*, and *ggplot2* packages.

**Figure 6.**
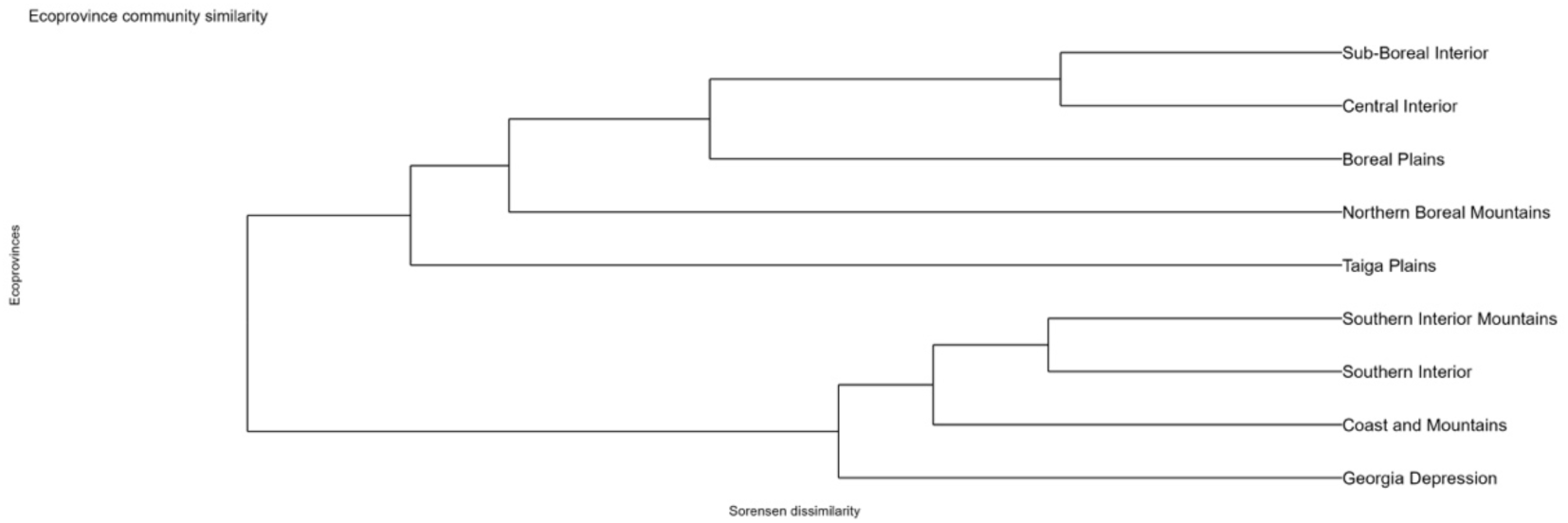
Dendrogram of Odonata community similarity for nine British Columbia ecoprovinces. Analysis is based on presence/absence data for each species, Sorenson dissimilarity, and UPGMA (unweighted pair group method with arithmetic mean) linking algorithm and was conducted in R (version 4.2.2) package *vegan*.

In addition to locality data, 19,234 records in the database (55.5%) included some degree of habitat description. These descriptors represented 1,567 unique descriptors of wetland habitats. Text mining of these unique descriptors revealed four categories of information (wetland type; abiotic factors; vegetation description; and size of habitat). These descriptive terms were present in a large number of combinations (Figure 7) with 206 (12.7%) of the descriptors including information in all four categories.

**Figure 7.**
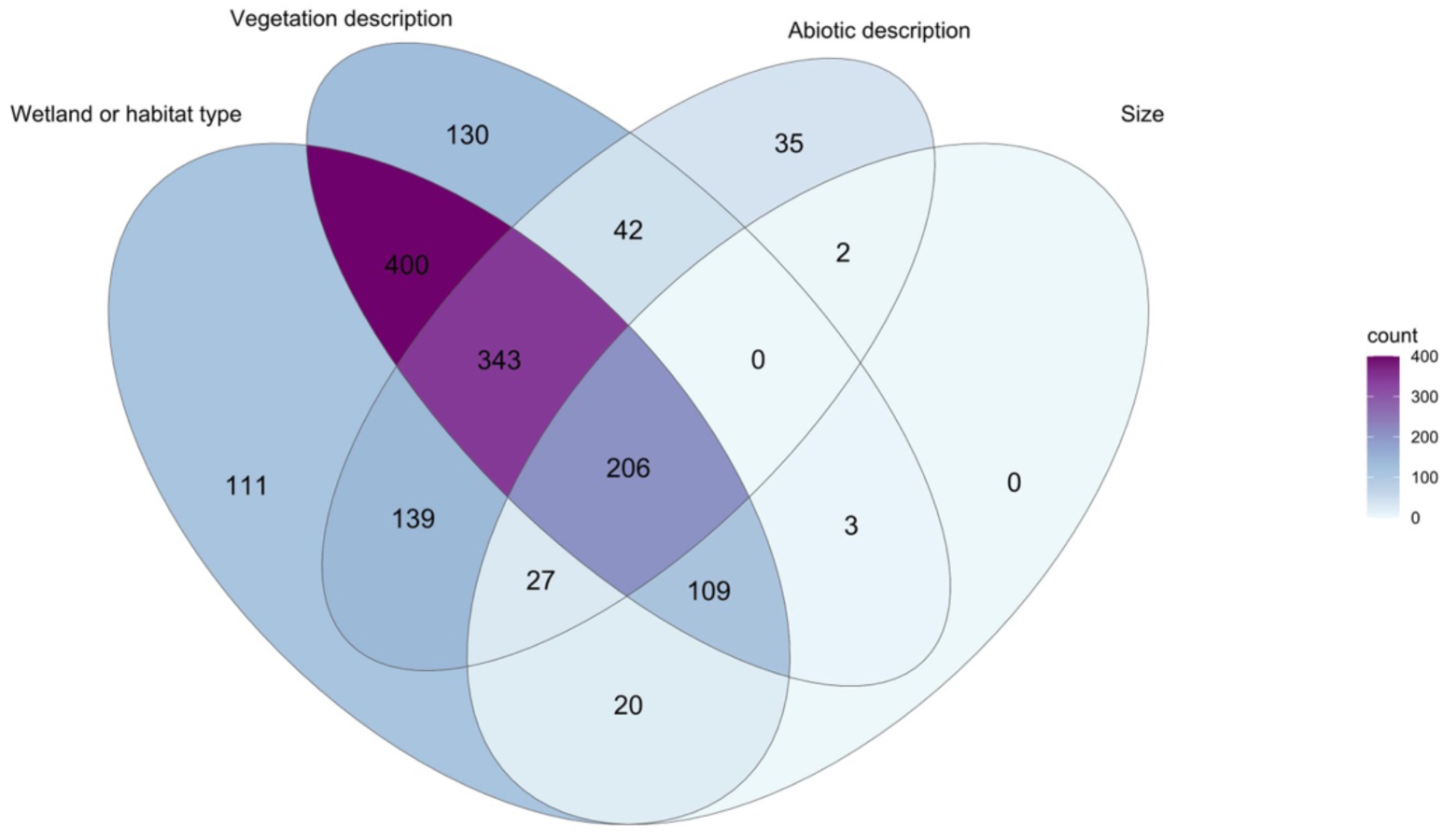
Venn diagram showing extent of overlap in specimen habitat descriptions for RBCM British Columbian Odonata. Diagram created in R version 4.2.3 with the packages *ggVenn* (Yan 2023) and *ggVenndiagram* (Gao 2022).

Of the 34,687 records in the database, 18,970 (54.7%) contained elevation data. However, by combining available latitude and longitude data and Google Earth Pro data, elevations could be estimated and included for an additional 10,548 records. This process allowed elevation comparisons to be made for 85.1% of the records. Figure 8 shows the percentage of records of each Odonata genus occurring in seven elevation strata (below 250m to above 3000m). Many genera included records at a range of elevations, but eight genera were restricted to below 500m and six genera had the majority of their records taken from above 750m. A pairwise analysis of the species similarity at different elevation strata revealed patterns that could provide guidance for conservation efforts as species shift range and elevation with accelerating climate change (Table 3). For instance, the species list for the 501m-750m elevation range was 92.1% similar to the species list for the 751m-999m elevation range. However, the species list for the 2000m-2999m elevation range was only 18.2% similar to the species list for the over 3000m elevation range. The similarity between the lowest elevation (below 250m) and the highest (over 3000m) species lists was very low (11.9%).

**Figure 8.**
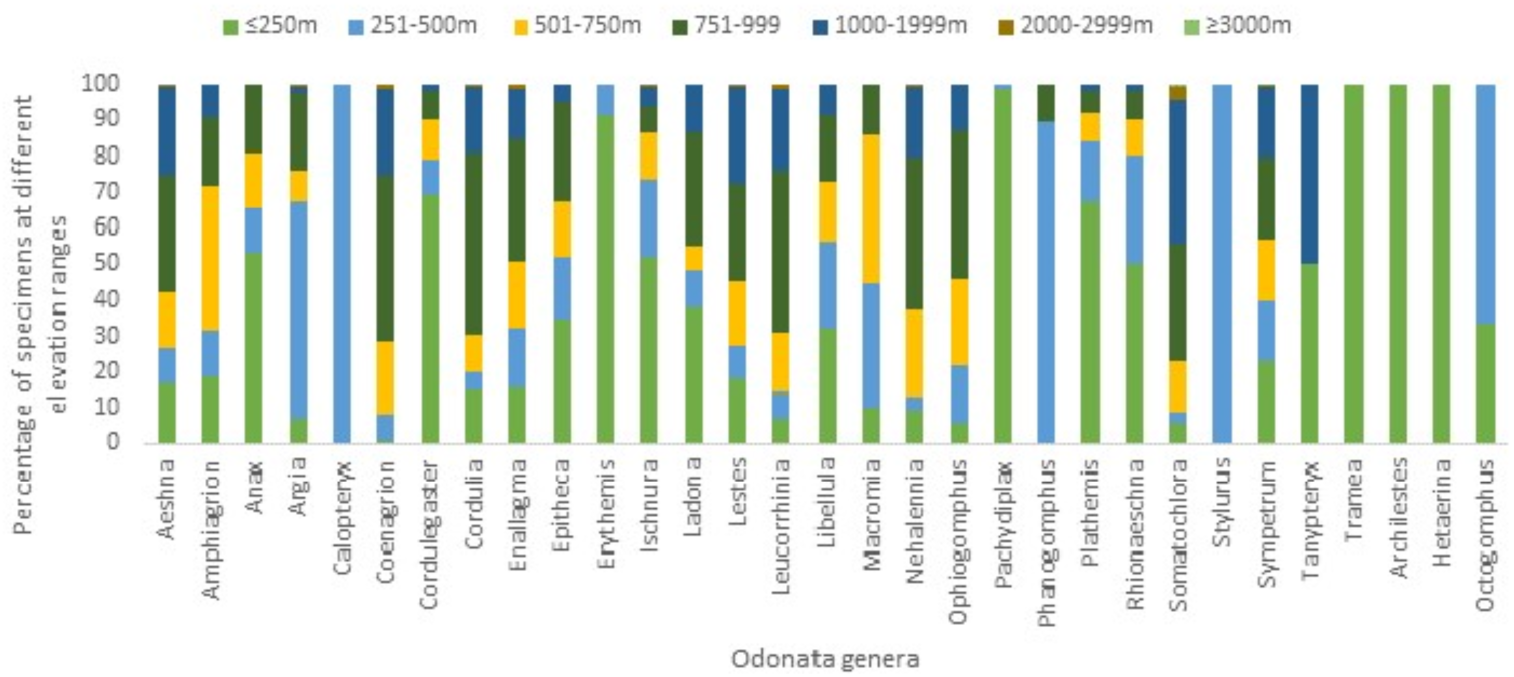
Percent composition of collected Odonata genera at each elevation range with both recorded and retrieved data. A total of 29,518 specimens were included, comprising 31 genera.

**Table 3.** Faunal similarity (FS) between elevation ranges (in m above sea level) using both recorded and retrieved elevation data. N_1_ – number of species in community 1. N_2_ – number of species in community 2. N_c_ – number of species in both communities. If elevation ranges have exactly the same species, FS should equal 1.

|  | <250:<br>251-500 | 251-500:<br>501-750 | 501-750:<br>751-999 | 751-999:<br>1000-1999 | 1000-1999:<br>2000-2999 | 2000-2999:<br>>3000 | <250: >3000 |
| --- | --- | --- | --- | --- | --- | --- | --- |
| $N_1$ | 66 | 76 | 74 | 72 | 64 | 30 | 66 |
| $N_2$ | 76 | 74 | 72 | 64 | 30 | 9 | 9 |
| $N_c$ | 59 | 66 | 70 | 62 | 29 | 6 | 8 |
| FS | 0.711 | 0.786 | 0.921 | 0.838 | 0.446 | 0.182 | 0.119 |

## Discussion

The RBCM Odonata collection digital database is a very complete, well- curated, and informationally rich biodiversity and conservation resource. With over 95% of records identified to species and over 81% of records including precise geographic data, it is as or more complete than many other entomology collections. For example, Favret and DeWalt (2002) found that, within the INHS collection, only 22% of Ephemeroptera records and 88% of Plecoptera records were identified to the species level following a concerted data digitization project. Following concerted digitization records for nearly twenty years, only 10% of the over 1.5 million specimens across all orders in the University of Alaska Museum Insect Collection had species identifications captured digitally (Sikes *et al*. 2017). An effort to digitize Odonata records from seven collections in the province of Quebec resulted in a list of 40,447 records, 91.3% of which were identified to the species level (Favret *et al*. 2020).

In their assessment of North American entomology collections, Cobb *et al*. (2019) noted that most (81%) of historic (pre-1965) entomological specimens are to be found in the largest North American collections. The RBCM Odonata collection is a counterexample to this with a large number of historic specimens in a relatively small collection (Figure 2) due to its regionally focussed nature. By choosing to target Odonata of a single Canadian province, the collectors, curators, and taxonomic experts created an expansive time-series of known, extant species from across the province. However, even with that effort, there are limitations in geographical representation (Figure 4). Likewise, a collection date analysis reveals that recent records are lacking for six red-listed species – and one species, *Enallagma clausum*, has not been collected for over a quarter of a century (Table 2). A lack of recent records for key species may indicate conservation concerns requiring updated assessments for some of those seven British Columbia Odonata species (Table 2). A decline in new, physical, North American records of Lepidoptera species in recent decades has been documented as well in other collections (Girardello *et al*. 2019). This gap in recent records may also be due to a shift amongst entomologists towards photographic vouchers and online records (*e.g.*, iNaturalist). Orders consisting mainly of large insects, such as Odonata, may be susceptible to this shift in data collection. Supplementing physical specimen databases with purely digital records may be beneficial – for instance by reducing the number of listed individuals taken from a limited population. But photographic collection also has biases and limitations depending on the temporal and spatial coverage of the data (DiCecco *et al*. 2021).

Entomological collection databases, even those focussing on a specific region, include gaps in geographical coverage. This fact can present a challenge when using collection data for analyses of species distributions and changes in those distributions due to anthropogenic effects (Kharouba *et* al. 2018). Looking at over 19 million digitized global Lepidoptera records, Girardello *et al*. (2019) found that spatial gaps persist in species distribution estimates, especially in areas not specifically designated for conservation and in areas with low density of roads. Once visualized (e.g. Figure 4), locality gaps can serve as a guide for future biomonitoring and conservation efforts. Detailed gap analyses, using geographic coordinates and species abundances, are used to asses the overall reliability of collection datasets for species distribution assessments (Ponder *et al*. 2001). Similar analyses are also used as a means to target survey work in unsampled areas or areas of potential conservation importance (Funk *et al*. 2005; Bini *et al*. 2006).

It is possible, using only the available digitized data, to combine existing data categories or other interacting databases to add further detail. Analyses that include species habitat preference similarities – involving factors such as elevation, latitudinal range, habitat type, and ecoprovince – are possible through post-extraction work (Figs. 5-8). Such meta-analyses allow testing of hypotheses of ecological interactions, historical biogeography, and human impact on the environment. A caveat is necessary, however, when the results of different analyses are considered together. For instance, since only 52.5% of Cordulegastridae records included full latitude and longitude data, the conclusions of subsequent analyses for this Family should be offered with less confidence than those for a Family with more complete geographical records.

An ecoprovince approach to analysing insect distribution in British Columbia has previously been used for neuropteroid orders (Scudder and Cannings 2009); conopid Diptera (Gibson 2017); and spheciform wasps (Ratzlaff 2015). In all three cases, a greater species richness was observed in the southernmost ecoprovinces compared to northern ecoprovinces. A similar pattern is shown here for Odonata (Figure 5). In all three previous studies, it was suggested that the distributional difference could be due to biogeographical history or due to a general lack of records from northern ecoprovinces. Our analysis also showed a separation in species similarity between the southern and coastal ecoprovinces and the north (Figure 6). It is expected that future climate change impacts will affect the abiotic conditions of ecoprovinces of British Columbia differently, with subsequent unique impacts on insect populations in those ecoprovinces (Haughian *et al*. 2012). Ecoprovince analysis of entomological collection records is useful because insect species are often not included in conservation and biodiversity policy decisions. The reasons for this exclusion include: a) a lack of data on species distribution; b) a lack of data on species abundance and changes in that abundance over time; and c) a lack of data on anthropogenic impacts on individual species (Cardoso *et al*. 2011). Digitized public entomological data can help to overcome all three of these limitations.

While not always included in entomological collections databases, elevation data can be added to specimen records *post hoc* through the use of other geographic databases. These data are highly valuable for downstream conservation and ecological analyses (Table 3). For example, high elevation aquatic insects are particularly vulnerable to future climate change due to abiotic factors (Birrell *et al*. 2020). Other research has found that low elevation populations of Lepidoptera have been more severely impacted by recent climate change than higher elevation populations (Halsch *et al*. 2021). Biotic factors, especially insect-plant interactions, are also likely to change dramatically along elevational gradients under future climate change effects (Rasmann, *et al*. 2014; Adedoja *et al*. 2020).

A bare minimum of adequate specimen preparation, complete labelling, identification to the family level, and digital capture is the goal for every specimen in an entomology collection (Favret et al. 2007). Records from old data sources can be used for new analyses, but when they are not already digitized it requires considerably more effort (Giberson and Burian 2017). This detailed analysis of the RBCM Odonata collection was only possible because of a previous concerted effort to digitally capture specimen data. This effort took place over a long period (1988-2001) and involved an investment of significant labour and funds (R.A. Cannings, *pers. comm.*). A similar analysis would not be as effective for other RBCM entomology collections due to a lack of digitization of all specimens and general data incompleteness. For example, 100% of the 272 drawers of Odonata specimens in the RBCM collection have been checked for specimen preparation completeness and entered into the database, whereas only 153 of the 394 drawers (38.8%) of Diptera specimens have received the same level of digitization. This means that a far smaller proportion of Diptera specimens are available for any sort of digital analysis. Furthermore, the proportion of complete records in digital databases is far lower in other taxa than it is for Odonata in the RBCM entomology collection. Only 20.7% of Diptera records, 36.1% of Coleoptera, 26.1% of Hemiptera, and 24.5% of Hymenoptera include species level identifications in the database.

An unexpected, but important, finding in these analyses was the relatively small number of trained experts whose work improves large datasets. Over thirty years ago it was noted that entomological collections suffer from “a critical lack of both talented young workers and secure jobs for them” (Miller 1991). The current study supports that assertion – only five taxonomists were responsible for nearly all of the species-level identifications in the British Columbia Odonata dataset (Figure 3). A lack of trained personnel to perform the necessary improvement and digitization of entomology collections would appear to be an imminent threat. Miller *et al*. (2020) emphasized the need to fund and maintain specific training in specimen preparation, data capture, and curation. These skills are not often included in the standard training of undergraduates and graduates not working directly in natural history collections. Miller (1991) noted that there are far too few programmes offering both museum experience and modern curatorial training.

New investments in entomology collection training and labour are absolutely necessary. Turney *et al*. (2015) emphasize that additional funding to natural history collections is necessary for them to meet their role as permanent storehouses of biological voucher data. The current rate of specimen digitization in North American entomology collections will need to increase by 400% to realistically capture most specimen data by the year 2050 (Cobb *et al*. 2019). This increased emphasis on biodiversity data capture and accessibility can, and should, be completed with an eye to spin-off research related to general conservation and societal needs (Cobb *et al*. 2019). That particularly includes conservation of biodiversity in the context of accelerating climate change. Nelson and Ellis (2018) list major governmental initiatives for digitally capturing and making available biodiversity data in the United States of America, Australia, Mexico, Brazil, Europe, and China. While these programs have proven successful elsewhere, a similar, well-funded government program does yet not exist for Canada.

## Conclusion

There are many public entomology collections in Canada. All of them are tasked with housing, organising, and making fully accessible, records of entomological biodiversity. Most collections focus on a specific region or province, although almost all contain specimens from other regions of Canada or beyond. Most collections have one or more taxonomic focus which can change over time with shifting priorities and personnel. None of these collections are private and all of their contained data could and should be made fully available to researchers in Canada and internationally. However, that effort will require funding for training and employing collections personnel.

This study was completed as a part of a graduate level course in entomological curation and the co-authors are the two instructors and five students of this course. The analyses presented were the product of a large-scale database extraction which was then analyzed by the five students with guidance from the instructors. The students did not have direct access to the collections and did not interact with the specimens. As such, this analysis truly represents what can be done remotely with entomology collections data, and how such analyses can inform biodiversity conservation. Discussions on taxonomic revision, georeferencing, and ecological notation were all components of the course. Those are just some of the skills necessary for those that wish to work with and improve entomological databases to explore ecological hypotheses and advise on conservation management policy.

Continued investment in training in entomological curation and collections management is essential. This will not only provide a key to increase the rate of digitally capturing the massive backlog of biodiversity data currently locked within collections, but it will also increase the quality and usefulness of existing digitized collections. With a new emphasis on training in curation and collection management skills at the undergraduate and postgraduate level, it will be possible to free even more biodiversity data from the locked cabinets of museums.

## Acknowledgements

We are very grateful to Dr. Rob Cannings, curator emeritus of entomology at the Royal BC Museum for his dedication to the Odonata collection and for his input on the writing of this manuscript.

